# Aurora A inhibition limits centrosome clustering and promotes mitotic catastrophe in cells with supernumerary centrosomes

**DOI:** 10.1101/401661

**Authors:** Bernat Navarro-Serer, Eva P Childers, Nicole M Hermance, Dayna Mercadante, Amity L Manning

## Abstract

The presence of supernumerary centrosomes is prevalent in cancer, where they promote the formation of transient multipolar mitotic spindles. Active clustering of supernumerary centrosomes enables the formation of a functional bipolar spindle that is competent to complete a bipolar division. Disruption of spindle pole clustering in cancer cells promotes multipolar division and generation of non-proliferative daughter cells with compromised viability. Hence molecular pathways required for spindle pole clustering in cells with supernumerary centrosomes, but dispensable in normal cells, are promising therapeutic targets. Here we demonstrate that Aurora A kinase activity is required for spindle pole clustering in cells with extra centrosomes. While cells with two centrosomes are ultimately able to build a bipolar spindle and proceed through a normal cell division in the presence of Aurora A inhibition, cells with supernumerary centrosomes form multipolar and disorganized spindles that are not competent for chromosome segregation. Instead, following a prolonged mitosis, these cells experience catastrophic divisions that result in grossly aneuploid, and non-proliferative daughter cells. Aurora A inhibition in a panel of Acute Myeloid Leukemia cancer cells has a similarly disparate impact on cells with supernumerary centrosomes, suggesting that centrosome number and spindle polarity may serve as predictive biomarkers for response to therapeutic approaches that target Aurora A kinase function.

## Introduction

The formation of a mitotic spindle with two spindle poles is integral to proper chromosome segregation. During mitosis a pair of centrosomes, the microtubule nucleating and organizing centers of the cell, serve as focal points at which molecular motors and crosslinker proteins cluster microtubules (reviewed in (1) and (2)). In cancer, centrosome amplification is common, where it contributes to aneuploidy, and is correlated with high tumor grade and poor patient prognosis (3-7). A direct consequence of having more than two centrosomes is the capacity to form a mitotic spindle with more than two spindle poles (8). Multipolar divisions that can result from such spindle geometry, lead to daughter cells with dramatic changes in chromosome number, and decreased viability (5, 9). To limit multipolar divisions, cells with excess centrosomes actively cluster them into two functional spindle poles (7, 10-13). Cells with clustered centrosomes complete bipolar anaphases with only moderate chromosome segregation errors, and remain viable (8, 14). In this way, the molecular players involved in centrosome clustering have an important role in determining the fate of cancer cells with supernumerary centrosomes and are therefore attractive candidates for therapeutic approaches (11, 15).

Aurora A (AurA) is a serine/threonine kinase that is important for proper bipolar spindle formation. Localized to the centrosome during interphase and to spindle poles during mitosis, AurA phosphorylates protein targets to regulate centrosome maturation and spindle assembly such that loss of AurA kinase–dependent phosphorylation of key mitotic targets disrupts spindle formation (16, 17). Some targets of AurA kinase, including Eg5, contribute to spindle formation by promoting anti-poleward forces that push centrosomes apart (18, 19). Other direct and indirect targets of AurA kinase, including TACC, PLK1, and NEK6, have roles in stabilizing microtubule dynamics and promoting spindle pole organization (20-23). Inhibition of AurA kinase activity is known to impair spindle pole organization in cells with two centrosomes (24-29). Because of its important mitotic role, AurA inhibitors are being explored as therapeutics to target rapidly growing cancer cells (30-32). However, clinical response to Aurora A inhibitors have been variable and it remains unclear which cancers may be most responsive to AurA inhibition.

Here we show that the response to AurA kinase inhibition is influenced by centrosome number and the capacity to form multipolar spindles during mitosis. As described previously, inhibition of AurA kinase activity in cells with normal centrosome content transiently disrupts mitotic spindle formation (24-29). However, these cells ultimately achieve spindle bipolarity to enable a bipolar division and sustained proliferation. In contrast, we show that AurA kinase activity is essential in cells with supernumerary centrosomes such that AurA inhibition compromises centrosome clustering, permits abortive cell division, and results in multinucleated, highly aneuploid daughter cells with reduced proliferative capacity. Together, these findings suggest that therapeutic approaches that exploit AurA inhibition will be most advantageous in cancers with supernumerary centrosomes.

## Results

### AurA promotes clustering of supernumerary centrosomes

To explore the relationship between spindle polarity and response to AurA inhibition we utilized two experimental systems to manipulate centrosome number. First, we induced supernumerary centrosomes using a previously well-characterized human epithelial cell line carrying a doxycycline-regulated PLK4 expression construct (indPLK4) (8). PLK4 is a regulator of centrosome biogenesis that, when overexpressed, leads to a cell-cycle dependent over-duplication of centrioles (Figure 1A & B). In the absence of PLK4 induction, the hTERT-immortalized human retinal pigment epithelial (RPE-1) cell line is diploid and contains a single pair of centrioles (one centrosome) during interphase. This centrosome is duplicated in preparation for mitosis, facilitating the formation of a bipolar mitotic spindle (Figure 1C & D). In the first cell cycle following induction of PLK4 overexpression cells generate excessive centrioles that remain associated as “rosettes”. In the second cell cycle following PLK4 induction, centriole disengagement and maturation results in supernumerary centrosomes. By 36 h post-PLK4 induction, more than 90% of interphase RPE-1 cells, and all observed mitotic cells have >4 centrioles, as indicated by centrin-2 staining (Figure 1A and B). Consistent with this increase in centriole number, following PLK4 induction, ∼60% of mitotic cells exhibit a multipolar spindle (Figure 1A-D, Supplemental Figure 1A & B). Nevertheless, by the time they proceed to anaphase, the vast majority (>80%) of indPLK4 cells have clustered excess centrosomes to form bipolar spindles (Figure 1D-F). A complementary approach to generate cells with extra centrosomes via cytokinesis failure was used to assess how centrosome number, independent of PLK4 over-expression, impacts spindle pole clustering. For this approach, the near-diploid HCT116 colon cancer cell line, which contains 2 centrosomes, was treated with Cytochalasin B (CytoB) to induce cytokinesis failure (Supplemental Figure 2). The resulting cells have double the DNA content, and double the centrosome number. Similar to the RPE-1 system described above, these cells also exhibit an increased incidence of multipolar spindles that resolve to cluster excess centrosomes for bipolar anaphases (Supplemental Figure 1B-F).

**Figure 1.**
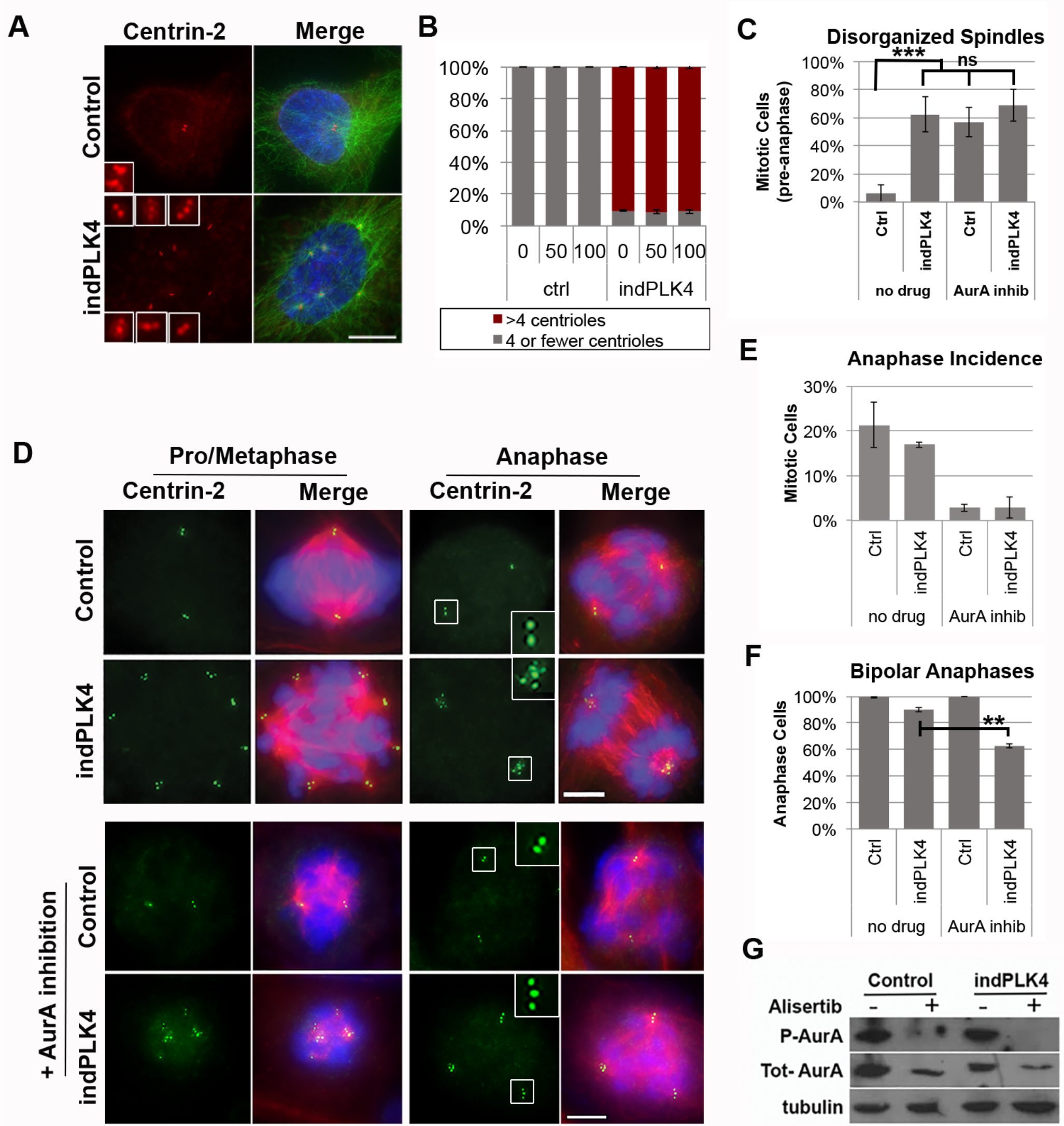
Aurora A kinase promotes clustering of supernumerary centrosomes. A & B) RPE-1 cells induced to overexpress PLK4 (indPLK4) have excess centrosomes. Insets show 4x enlargements of individual centrioles (centrin-2 in red) within a single cell. C-F) Following inhibition of Aurora A kinase activity with 100nM alisertib, cells with two or excess centrosomes similarly exhibit disorganized mitotic spindles. Cells with extra centrosomes are efficiently clustered into bipolar spindles prior to anaphase onset while those with supernumerary centrosomes undergo multipolar mitoses. Centrin-2, a marker of centrioles is shown in red (panel A) or green (panel D), tubulin in red, and chromatin in blue. The scale bars are 10 μM. Error bars are SEM, * = p < 0.05, **= p < 0.01, ***= p < 0.001.

Alisertib (MLN8237) is an orally bioavailable inhibitor of AurA kinase that is ∼200 fold more selective for AurA than the closely related Aurora B (33). Pharmacological inhibition of AurA kinase activity can be monitored through loss of AurA auto-phosphorylation of threonine residue 288 in its activation loop (34). Within 2 hours, 100nM alisertib is sufficient to inhibit AurA kinase activity and prevent threonine 288 phosphorylation (p- AurA) in mitotic cells, irrespective of centrosome number (Figure 1G).

To assess how mitotic cells with excess centrosomes respond to AurA inhibition, both control and indPLK4 RPE-1 cells, and HCT116 cells +/- Cyto B were treated with inhibitor for 16 hours, followed by immunofluorescence imaging. This duration of treatment was sufficient to limit each cell to one mitotic event in the presence of AurA inhibition. Consistent with previous reports, we find that cells with two centrosomes exhibit an increase in acentrosomal and disorganized mitotic spindle poles following exposure to any one of four specific inhibitors of AurA kinase activity: alisertib, MLN8054 (MLN), Aurora A inhibitor 1 (AA1), and MK-5108 (MK/VX-689) (Supplemental Figure 1B) (22). Nevertheless, nearly all anaphase and telophase cells in these populations were bipolar (Figure 1D & F, Supplemental Figure 2C-F), indicating that even in the context of AurA inhibition acentrosomal spindle poles are eventually focused and spindle bipolarity is achieved prior to anaphase onset.

Following Aurora A inhibition, cells with supernumerary centrosomes form multipolar and disorganized spindles similarly to control cells. In these cells centrosomes are present at the majority of excess spindle poles (Figure 1C and D) and there is a significant decrease in the proportion of anaphase cells with bipolar spindles (Figure 1D and F, Supplemental Figure 2C, D & F). Together, this data suggests that cells with extra centrosomes are unable to achieve sustained centrosome clustering.

### Cell fate in the presence of AurA inhibition is influenced by centrosome number

Cells that are unable to form a bipolar spindle are expected to accumulate in mitosis. However, FACs analysis of cellular DNA content, together with imaging-based assessment of mitotic enrichment indicate that the 4N (G2/M) population of cells is not significantly changed and mitotic cells do not surpass ∼10% of the cell population following short term (16-24 h) AurA inhibition (Supplemental Figure 1C & D). Together, this suggests that mitotic defects imposed by AurA inhibition are either transient, or lethal for cells with excess centrosomes. To differentiate between these two possibilities, we performed live cell imaging of control cells, and those with supernumerary centrosomes in the presence or absence of AurA inhibition.

Aurora A is known to function in both centrosome maturation and spindle assembly pathways and long term inhibition or RNAi-based depletion strategies compromise both processes. Therefore, to assess the role of AurA specifically in spindle bipolarity in cells with excess centrosomes, while limiting confounding effects of AurA inhibition on centrosome maturation, we performed live cell imaging on cells that entered mitosis within the first 30 minutes of drug-induced AurA inhibition (ie after centrosome maturation). These cells were then followed though mitotic exit and for the next 36 hours to assess cell fate. Images of RPE-1 cells expressing an RFP-tagged histone (RFP-H2B) to enable monitoring of chromosome movement and cell cycle progression were captured every 5 minutes for the duration of the experiment. Untreated RPE-1 cells progressed from nuclear envelope breakdown (NEB) to anaphase onset in ∼20 minutes. Consistent with our fixed cell analysis, following a prolonged mitosis ∼85% of cells with extra centrosomes (indPLK4) ultimately exit mitosis with a bipolar division (Figure 2C), while the remainder complete a multipolar division (Figure 2A-C: indPLK4).

**Figure 2:**
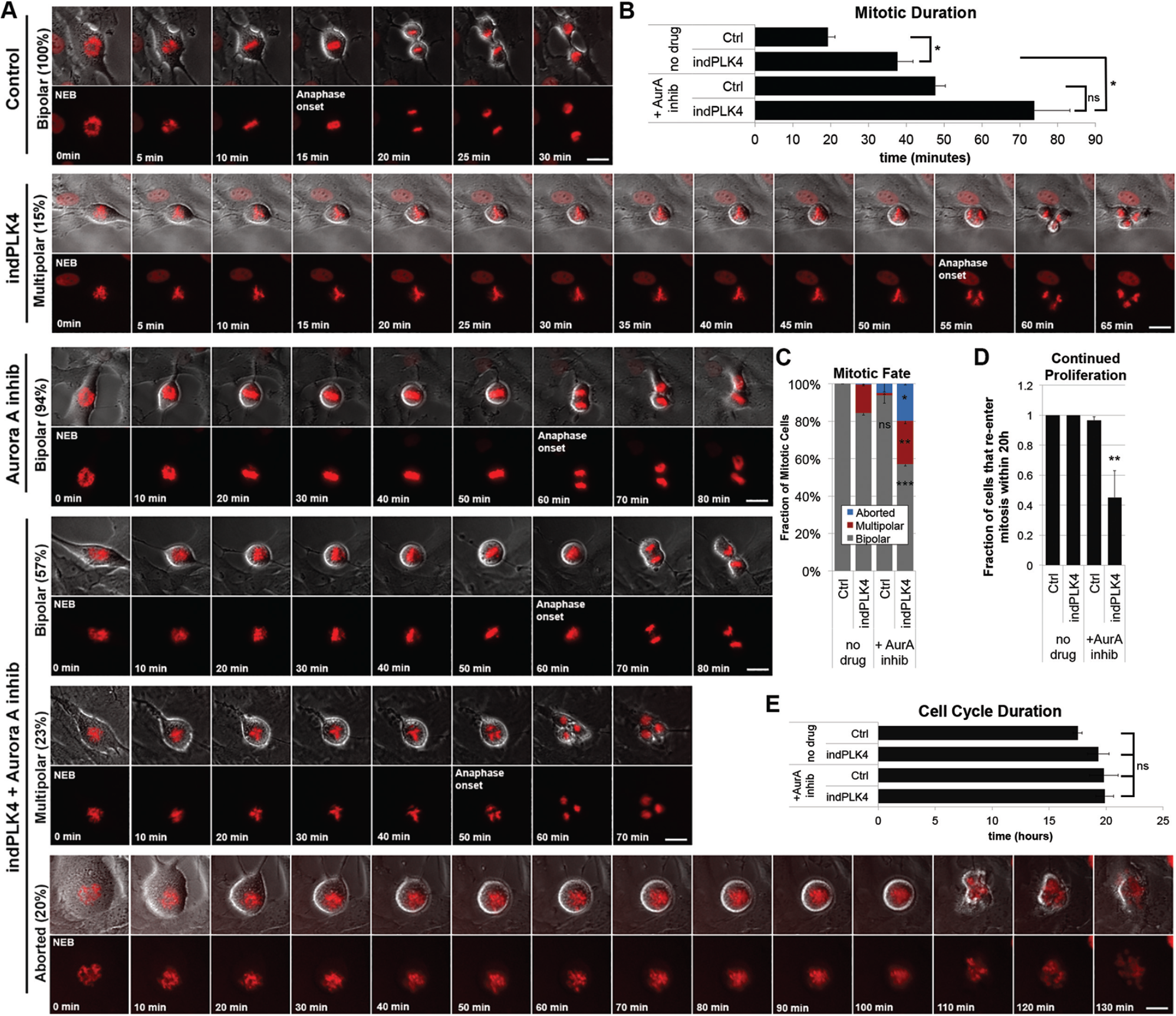
Centrosome number influences mitotic outcome following Aurora A kinase inhibition. A) Representative still frames from live cell phase contrast and fluorescence imaging of RFP-labeled Histone 2B (in red)-expressing cells. Nuclear envelope breakdown (NEB) to anaphase onset and cytokinesis are shown. Note that 10 min increments are represented for cells treated with Aurora A inhibition, 5 min increments for those not treated. Scale bars are 10μM. B) The presence of supernumerary centrosomes promotes an increase in the time from NEB to anaphase onset following Aurora A kinase inhibition. C) Both control cells and those with PLK4 induction alone divide to form two daughter cells following metaphase alignment of chromosomes. In the presence of Aurora A kinase inhibition, the majority of control cells also complete metaphase alignment and divide into two daughter cells. However, cells with PLK4 induction often attempt cytokinesis in the absence of chromosome alignmnet. D) In the presence of Aurora A inhibition, cells with PLK4 induction are less likely than cells without Aurora A inhibition and/or without PLK4 induction to proceed to a second mitotic division within 20 hours of a first division. E) Cells in all conditions that do complete a bipolar division continue to a second division with comparable cell cycle timing. Error bars are SEM, * = p < 0.05, **= p < 0.01, ***= p < 0.001.

Consistent with a high incidence of bipolar anaphases seen in fixed cell imaging (Figure 1C-F), following AurA inhibition control cells persist through a prolonged mitosis to achieve full alignment of chromosomes along the cell equator and complete a bipolar division (Figure 2A, B & C). indPLK4 cells similarly exhibit a mitotic delay, but live cell imaging revealed that only half are competent to complete a bipolar division in the presence of AurA inhibition. For the cells that are unable to complete a bipolar division in the presence of AurA inhibition, mitotic exit is catastrophic. In agreement with the high incidence of multipolar spindles seen in fixed cell analysis of anaphase (Figure 1), ∼20% of cells with supernumerary centrosomes achieve chromosome alignment and exit mitosis via a multipolar or abortive division. The remaining 20% of cells attempt cytokinesis in the absence of metaphase alignment or obvious anaphase segregation (Figure 2A, B & C).

Following completion of mitosis, each cell was tracked to monitor cell cycle re-entry. indPLK4 cells that were able to achieve a bipolar division were just as likely as their counterparts with normal centrosome content to proceed through a subsequent cell cycle and enter into a second mitosis (Figure 2D & 2E: ∼90% enter a second mitosis, with an average cell cycle ∼19 hours). Consistent with previous reports (8, 14), we find the progeny of multipolar divisions exhibit decreased proliferative capacity and decreased viability such that none of the cells that experienced multipolar or abortive mitotic divisions were competent to re-enter the cell cycle within the duration of our movies. This failure to re-enter the cell cycle contributes to a 40% decrease in proliferation of cells with supernumerary centrosomes following AurA inhibition (Figure 2D).

Consistent with the high incidence of abnormal and abortive anaphases, following a single cell cycle in the presence of AurA inhibition, immunofluorescence analysis demonstrates that nearly 25% of indPLK4 these cells are multinucleated, a significant increase over that seen in either indPLK4 cells alone, or in control cells following AurA inhibition (Figure 3 A & B). Fluorescence *in situ* hybridization (FISH)- based approaches using centromere-targeted enumeration probes for individual chromosomes similarly demonstrate that while short term induction of PLK4 (<2 cell cycles) results in a 2-fold increase in aneuploid cells, indPLK4 cell populations exhibit a 4-fold increase in aneuploid cells and a >6-fold increase in tetraploid cells following AurA inhibition (Figure 3C & D).

**Figure 3:**
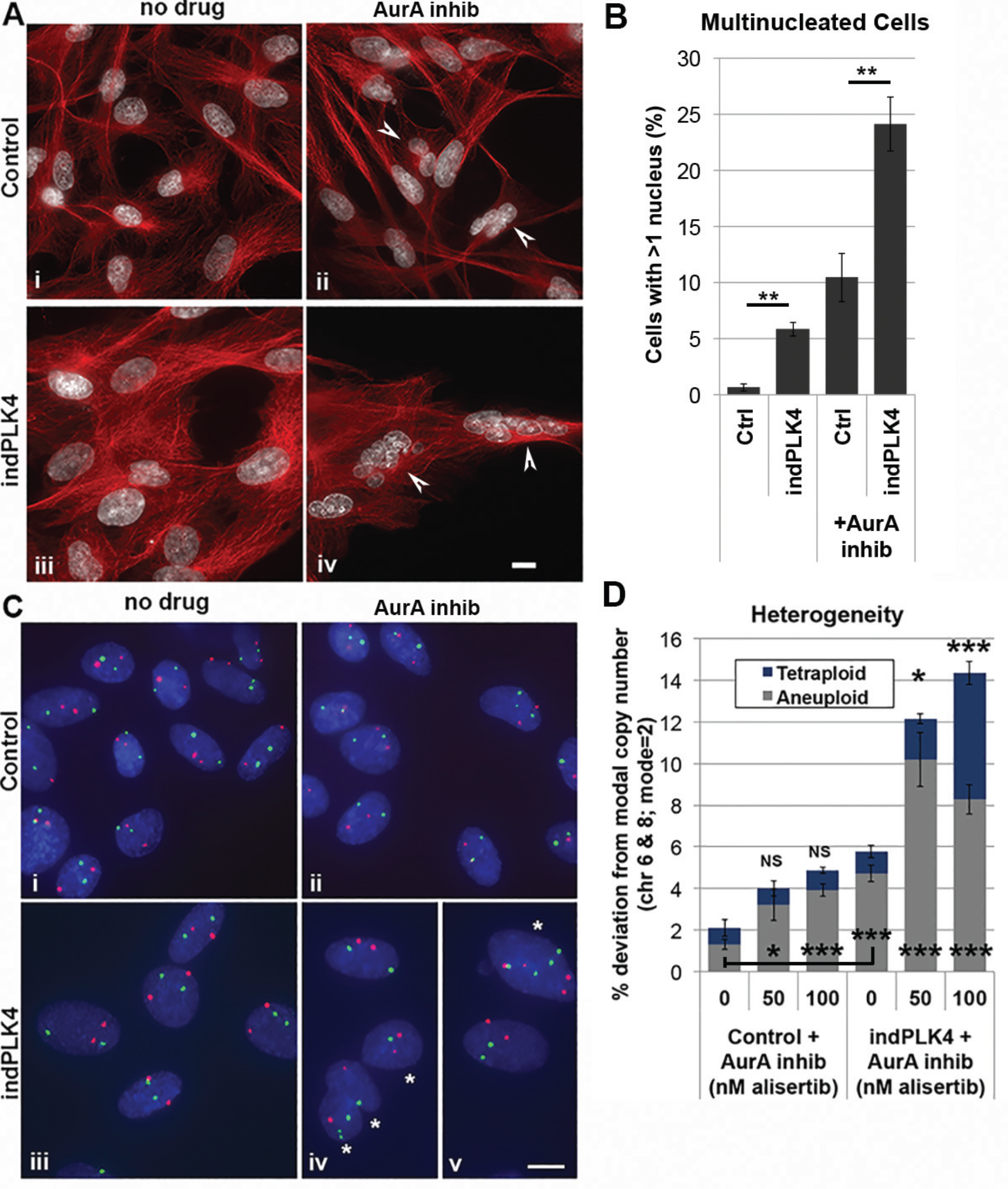
Aurora A inhibition compromises genome stability in cells with supernumerary centrosomes. A & B) Cells with supernumerary centrosomes exhibit high levels of multinucleated and C & D) tetraploid cells following inhibition of Aurora A kinase activity. A) white arrow heads indicate multinucleated cells. Tubulin shown in red, chromatin in white. C) Stars in iv and v indicate a multinucleated and mononucleated tetraploid cell, respectively. Probe for chromosome 6 shown in red and chromosome 8 shown in green. Scale bars are 10μM. Error bars are SEM, * = p < 0.05, **= p < 0.01, ***= p < 0.001.

### Centrosome number corresponds with response to Aurora A inhibition

To determine if centrosome number similarly corresponds with response to AurA inhibition in cancer cells we utilized a panel of Acute Myeloid Leukemia (AML) cell lines. AML is a cancer context where AurA kinase is often amplified, and where its inhibition is actively being exploited in preclinical and clinical approaches (35-37). Western blot and qPCR analyses demonstrate that AurA expression in these cell lines, independent of cell cycle distribution, range from levels comparable to that of the non-transformed RPE-1 cell line (ie HL60), to levels over 8-fold higher than that seen in RPE-1 cells (ie K562) (Figure 4A & B). Importantly, irrespective of AurA protein level, exposure to 100nM alisertib was sufficient to completely inhibit AurA kinase activity in each cell line, as judged by AurA auto-phosphorylation (Figure 4B), and resulted in enrichment of G2/M cells (Supplemental Figure 3).

**Figure 4.**
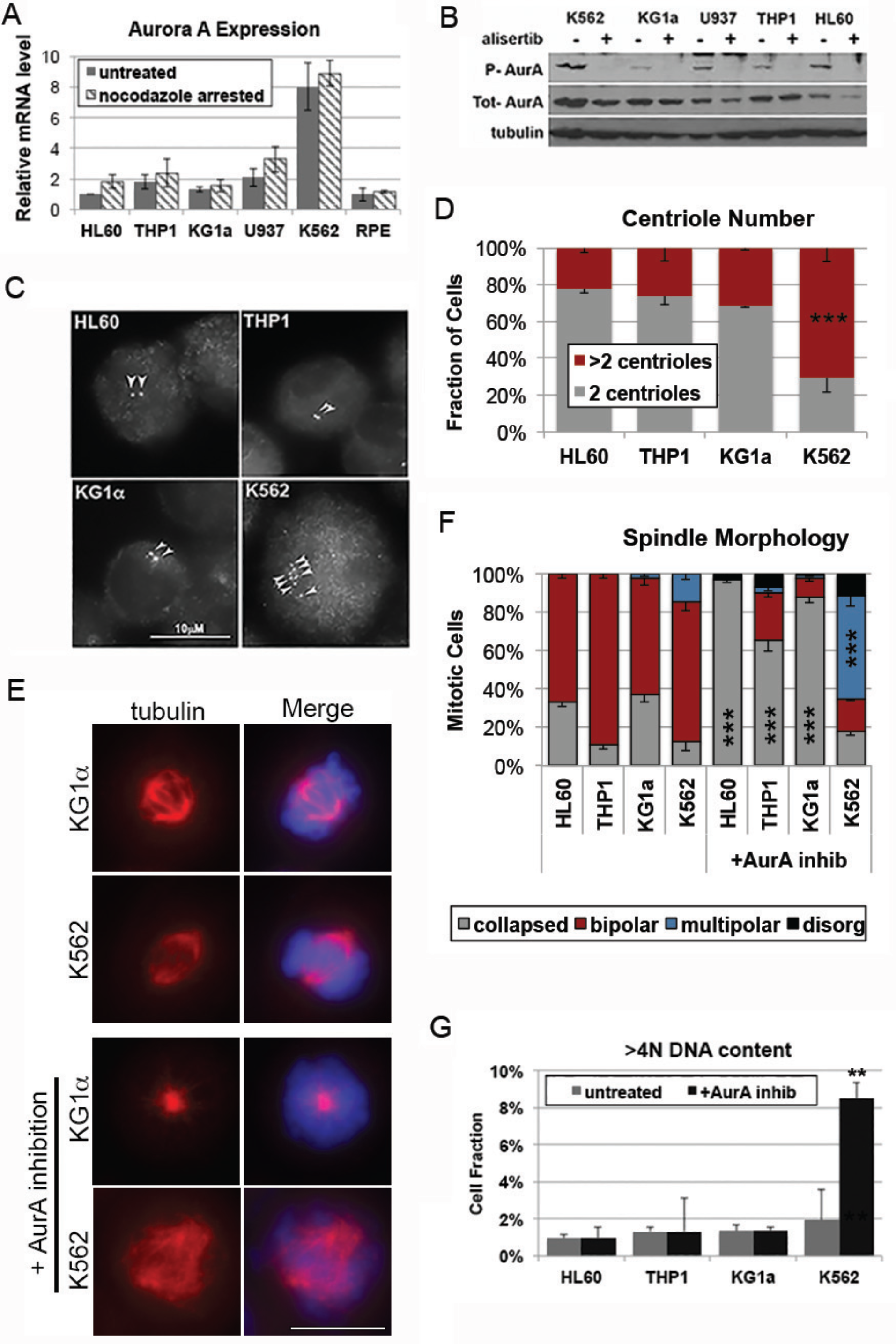
Response to Aurora A inhibition corresponds with centrosome number in Acute Myeloid Leukemia cells. A) A panel of AML cell lines exhibit a range of Aurora A expression levels, irrespective of differences in cell cycle distribution or proliferation between cell lines. B) Treatment with 100nM alisertib in all five cell lines is similarly sufficient to suppress auto-phosphorylation of Aurora A and promote enrichment of cell in G2/M (see also Supplemental Figure 3). C & D) Immunofluorescence analysis of centrin-2 staining indicates that K562 cells have supernumerary centrosomes. E & F) Following Aurora A inhibition, a significant increase in multipolar spindles is seen in K562 cells, which have supernumerary centrosomes, but not in cell lines with two centrosomes. G) FACs analysis of DNA content indicates that cells with supernumerary centrosomes also exhibit an increase in DNA content following Aurora A inhibition. Error bars are SEM, * = p < 0.05, **= p < 0.01, ***= p < 0.001.

While FISH-based analysis of chromosome copy number indicates that all 4 AML cell lines have a modal chromosome copy number near 4 (Supplemental Figure 4), immunofluorescence analysis of centrin-2 in these cell lines indicated that only the K562 cell line contains a significant portion of cells with >2 centrosomes (2 centrioles/centrosome) (Figure 4C & D). Nevertheless, mitotic spindles in each cell line are predominantly bipolar, indicating that K562 cells can efficiently cluster supernumerary centrosomes (Figure 4E & F). As seen with the RPE-1 and HCT116 cells, inhibition of AurA initially disrupts spindle bipolarity in all AML cell lines, regardless of centrosome number (Figure 4E & F). However, FACs and FISH-based analysis demonstrate cellular ploidy remains largely unchanged in AML cells with two centrosomes when AurA is inhibited, suggesting these cells are ultimately able to achieve spindle bipolarity and complete a bipolar division. In contrast, K562 cells, which have supernumerary centrosomes, experience mitotic failure in the presence of AurA inhibition, resulting in an increase in DNA content (Figure 4G).

## Discussion

Our data support a model whereby the presence of extra centrosomes, acquired either through over-duplication or a previous failed cytokinesis, can alter the response of cells to inhibitors of mitotic spindle assembly and subsequently impact the efficacy of such drugs. We find that following AurA inhibition, cells with supernumerary centrosomes exhibit multipolar and disorganized spindles that cannot be resolved prior to mitotic exit. Instead, such cells are subject to abortive mitoses where cytokinesis is attempted in the absence of metaphase alignment, and anaphase chromosome segregation is lacking. Daughter cells that result from such catastrophic mitoses are multinucleated and exhibit dramatic changes in chromosome content. Importantly, such progeny are not competent for continued proliferation. In contrast, following AurA inhibition, cells with normal centrosome number ultimately overcome a transient disruption of spindle geometry to form bipolar spindles and complete near-normal cell division. Progeny from these divisions remain proliferative and proceed through the cell cycle to enter a second mitosis with frequency and timing comparable to control cells. In this way, we propose that a differential response to AurA inhibition in cancer cells may stem not from a change in the initial impact on spindle assembly, but instead from an essential and cancer-specific dependence on AurA-driven dynamics of spindle-pole clustering activity in cells with supernumerary centrosomes.

Proteins that are required for centrosome clustering are often overexpressed in human cancers with extra centrosomes (reviewed in: (38)). Indeed, supernumerary centrosomes, and the need to cluster them to achieve proper cell division, has been proposed to be a driving force in the selective overexpression of proteins with pole-clustering functions. Consistent with this model, we find that AurA expression scales with centrosome number in our panel of AML cell lines (Figure 4A & E). Furthermore, AurA has been reported to be amplified or overexpressed in a number of cancer contexts, including colon, breast, and hematological cancers (39-43), where its expression corresponds with centrosome amplification (44-47).

Together with cell biological evidence that inhibition of AurA compromises mitotic progression, an increased AurA expression profile in cancer supports that AurA is a clinically relevant drug target (48). Pharmaceutical companies have begun to exploit this possibility, bringing numerous targeted inhibitors to the market as anti-cancer drugs, and spurring clinical trials (37, 49-55). However, while the efficacy and safety of inhibiting AurA have been supported by preclinical and early stage clinical trials, response to AurA inhibition has been limited (37, 55-57) and it remains unclear which patients may best benefit from AurA inhibitor treatment either alone, or in combination with standard chemotherapeutic approaches. We now show that cells with extra centrosomes are susceptible to catastrophic cell divisions when AurA activity is inhibited. Furthermore, consistent with the documented tolerance of patients to AurA inhibitors in the clinic (37, 56, 58), our data demonstrate that normal cells are able to overcome spindle disruption following AurA inhibition. Cells with two centrosomes can ultimately focus acentrosomal spindle poles to achieve bipolarity and exit mitosis with a bipolar anaphase. These divisions are preceded by chromosome alignment, occur with near normal mitotic timing, and resulting progeny remain proliferative. Together, these findings suggest that AurA inhibition can selectively corrupt proliferation of cancer cells with extra centrosomes, and that clinical assessment of mitotic structures may aid in identifying patients most likely to benefit from their use.

## Materials and Methods

### Cell culture, induction, and inhibition

Cells were grown in DMEM/F12 medium (RPE-1) or RPMI 1640 medium (HL60, K562, KG1α, and THP1) supplemented with 10% fetal bovine serum (FBS) and 1% penicillin/streptomycin and maintained at 37°C with 5% CO_2_. For experiments where PLK4 expression was induced, media was supplemented with 2 μg/mL Doxycycline for 36 hours. Alternatively, cells were treated with 30 μM Cytochalasin B for 16 h to induce cytokinesis failure and generate tetraploid cells with double the normal centrosome number (8). Drug treatments to subsequently inhibit AurA kinase (alisertib; MLN8054, Aurora A inhibitor 1, MK-5108 (VX-689): Selleckchem) were performed at the indicated concentrations for 16- 18 hours for immunofluorescence and FACs analysis, or as otherwise indicated. Unless otherwise noted, all experiments to inhibit Aurora A kinase were performed with the highly specific Aurora A kinase inhibitor alisertib at a concentration of 100nM.

### FACS and expression analysis

To assess cell cycle differences and response to mitotic inhibitors, cells were treated with 100 ng/mL Nocodazole (to induce mitotic arrest) and/or 100 nM alisertib for 16-18 hours, fixed and stained with propidium iodide, and processed for FACs analysis as previously described (59). To assess AurA expression levels, RNA extraction was performed with Qiagen’s RNeasy kit, and qRT PCR with Applied Biosystem’s SYBR green kit according to the respective manufacturers’ guidelines. Primers used for amplification of GAPDH and AurA were 5’-ccctctggtggtggcccctt-3’ & 5’-ggcgcccagacacccaatcc −3’ and 5’- ttttgtaggtctcttggtatgtg-3’ & 5’-gctggagagcttaaaattgcag-3’ respectively. Inhibition of AurA kinase activity was confirmed by western blot analysis of total AurA (Cell Signaling) and phosphorylated AurA levels (Cell Signaling), using tubulin as a loading control (dm1α, Sigma).

### FISH, immunofluorescence and live cell imaging

Cells were prepared and fixed as described previously (59) and α-satellite probes for chromosomes 6 and 8 hybridized to chromatin. Numerical Heterogeneity (NH) was determined by scoring a minimum of 500 cells per population for copy number of each chromosome probe, for each of three independent experiments. Percent variation from the modal copy number was calculated and compared across conditions.

For immunofluorescence imaging, cells were cultured on glass coverslips (RPE-1), or spun onto coverslips by centrifugation at 1000 RPM for 3 minutes (AML cell lines) prior to fixation for 10 min in ice-cold methanol, blocked in TBS-BSA (10 mM Tris at pH 7.5, 150 mM NaCl, 1% bovine serum albumin), and stained for α-tubulin (dm1α, Sigma), and/or centrin-2 (Santa Cruz) as indicated. DNA was detected with 0.2μg/mL DAPI (Sigma), and coverslips mounted with Prolong Antifade Gold mounting medium (Molecular Probes). Cells were plated in 12 well imaging bottom tissue culture dishes for live cell imaging. Images of fixed and live cells were captured with a Zyla sCMOS camera mounted on a Nikon Ti-E microscope with a 60x Plan Apo oil immersion objective, or a 20x CFI Plan Fluor objective. A minimum of 50 mitotic cells or 1000 interphase cells were analyzed per condition for each of 3 biological replicates. Live cell images were captured at 10 coordinates per condition at 5 min intervals for 24 hours, for each of 3 biological replicates. 100 individual cells were tracked through mitosis and analyzed for each condition and replicate. Error bars throughout represent standard error unless otherwise indicated. Pairwise analyses were performed using a one-way ANOVA to determine statistical significance.

## Acknowledgements

The authors would like to thank Neil Ganem for sharing cell lines, and Margaret Pruitt, Elizabeth Crowley, and Lily Kabeche for critical reading of the manuscript. ALM is supported by funding from NIH grant R00 CA182731 and the Smith Family Award for Excellence in Biomedical Research.

