## Supplemental Figures for "Aurora A inhibition limits centrosome clustering and promotes mitotic catastrophe in cells with supernumerary centrosomes"

**Supplementary Figure 1: Short-term inhibition of Aurora A disrupts spindle structure without altering overall cell cycle distribution.** ) PLK4 expression is efficiently induced for 36h from a doxycycline-regulated expression construct in RPE-1 cells to promote centriole over-duplication (see also figure 1). B) A panel of four clinically relevant, Aurora A inhibitors (alisertib; MLN8054, Aurora A inhibitor 1, and MK-5108 (VX-689)) all similarly corrupted spindle structure in RPE-1 cells with or without supernumerary centrosomes (indPLK4). C and D) Inhibition of Aurora A kinase activity with 100nM alisertib for 16h does not alter cell cycle distribution as judged by FACs analysis of DNA content, and only moderately increases the fraction of cells observed to be in mitosis by immunofluorescence based assay.

**Supplementary Figure 2: Inhibition of supernumerary centrosomes clustering is a general result of Aurora A inhibition and not specific to treatment with alisertib**. A) Treatment of HCT116 p53-/- cells with cytochalasin B (Cyto B) to induce cytokinesis failure results in binucleated cells with extra centrosomes. B) Similar to indPLK4 expressing RPE cells, HCT116 cells treated with Cyto B are able to efficiently cluster extra centrosomes to achieve a bipolar spindle prior to anaphase. C and D) HCT116 cells with and without extra centrosomes form multipolar spindles when Aurora A is inhibited. C-F) In the presence of AurA inhibition, cells with 2 centrosomes are able to achieve spindle bipolarity prior to anaphase onset, while cells with extra centrosomes primarily exit mitosis with multipolar spindles.

**Supplemental Figure 3: Cell cycle response to Aurora A inhibition is similar in AML cells with two or supernumerary centrosomes.** A) FACs analysis indicates that all AML cell lines exhibit an initial enrichment in cells with 4N (G2/M) DNA content following short term exposure to the Aurora A inhibitor alisertib. C) Immunofluorescence analysis of mitotic figures shows that Aurora A inhibition induces a concentration-dependent enrichment in mitotic cells similarly in all four of these cell lines, while only promoting multipolar spindle structures in the cell line with supernumerary centrosomes (see also Figure 4F).

**Supplemental Figure 4: Ploidy does not correspond with centrosome number in AML cell lines.** FISH-based analysis of copy number for chromosomes 6 (green) and 8 (red) indicate that each of the 4 AML cell lines investigated had a modal chromosome copy number near 4, while only K562 cells exhibited supernumerary centrosomes (see Figure 4), indicating that ploidy alone is not a strong indicator of centrosome number.

Supplemental Figure 1

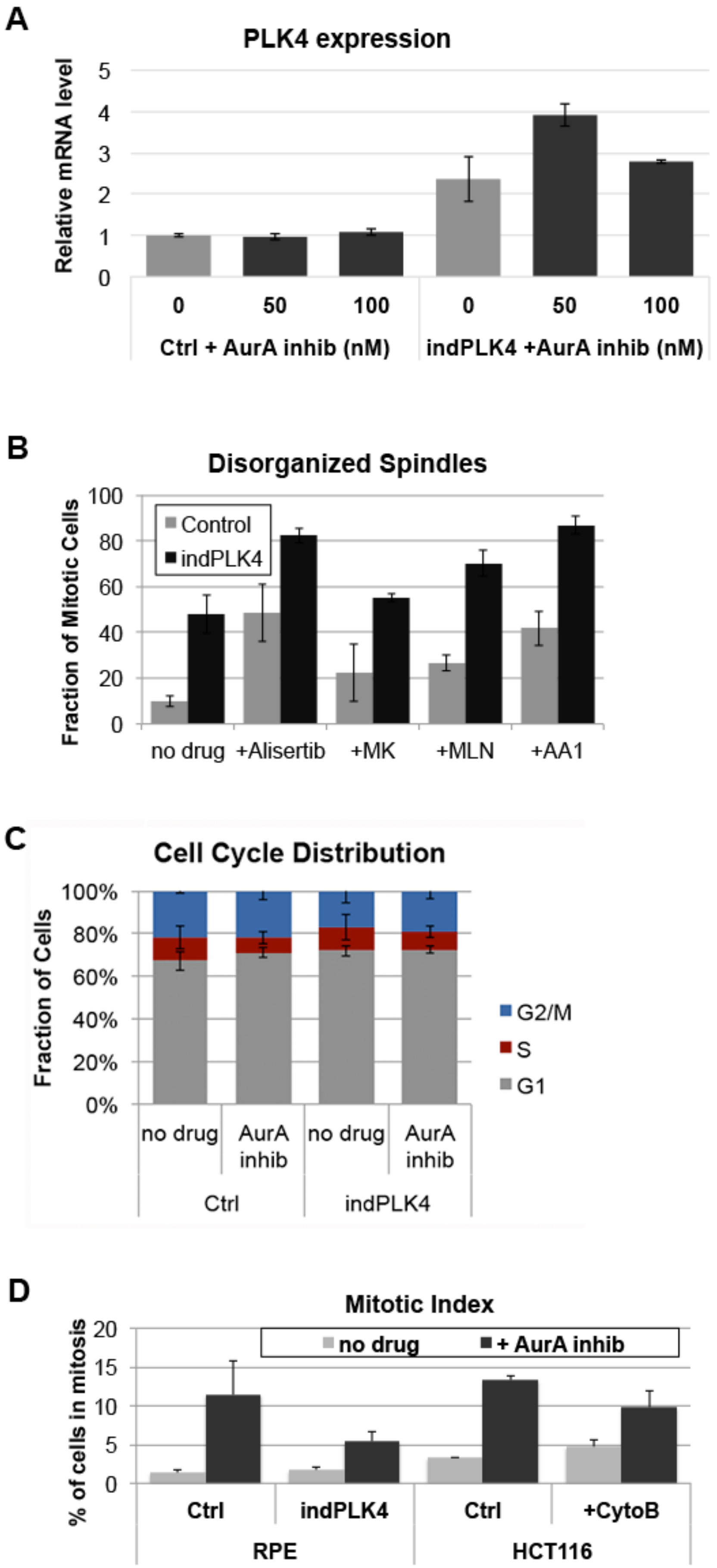

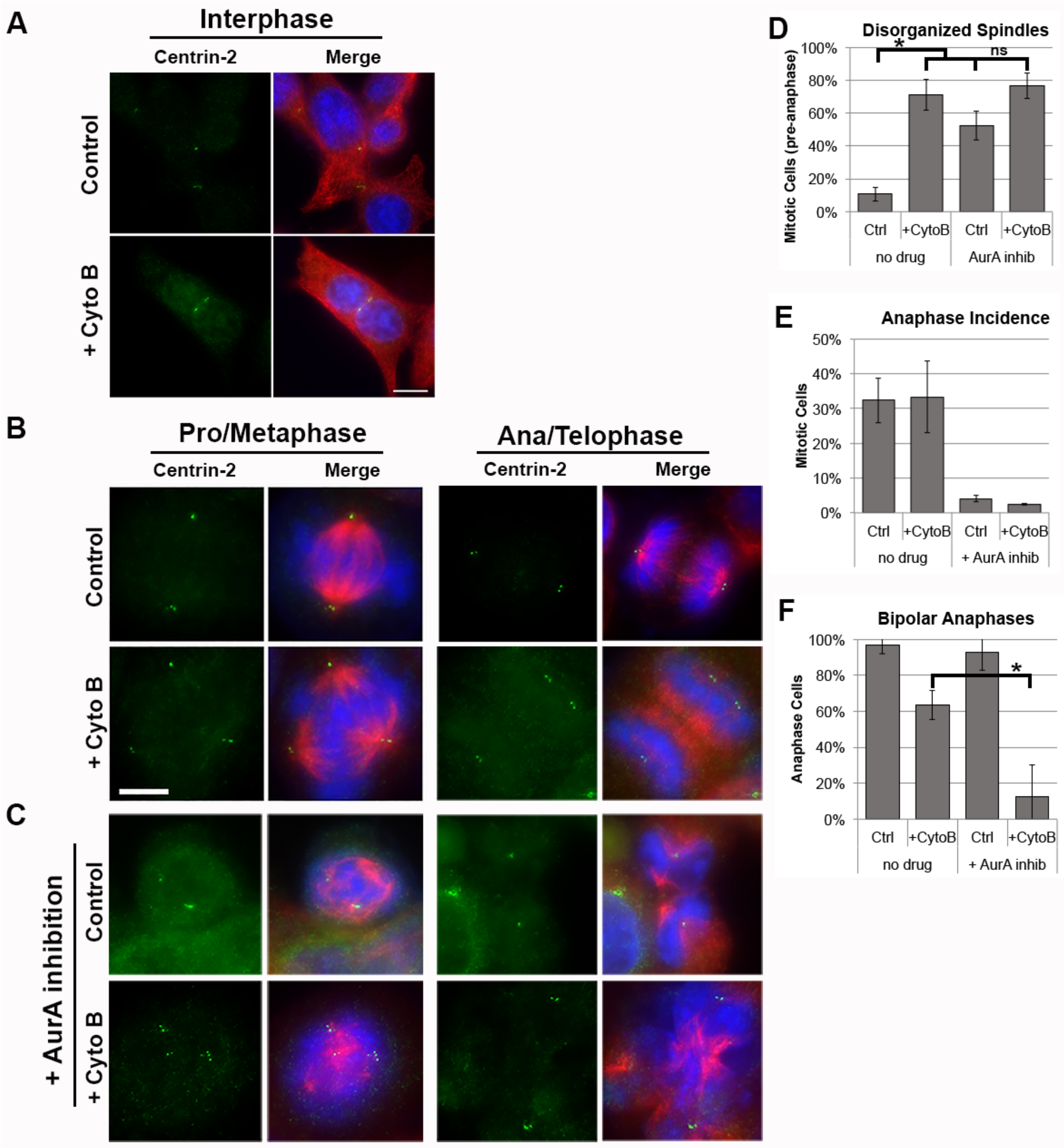
A
Interphase
Centrin-2
Merge
Control
+ Cyto B
B
Pro/Metaphase
Ana/Telophase
C
+ AurA inhibition
D
Disorganized Spindles
Mitotic Cells (pre-anaphase)
100%
80%
60%
40%
20%
0%
*
ns
Ctrl
+CytoB
no drug
AurA inhib
E
Anaphase Incidence
Mitotic Cells
50%
40%
30%
20%
10%
0%
+ AurA inhib
F
Bipolar Anaphases
Anaphase Cells

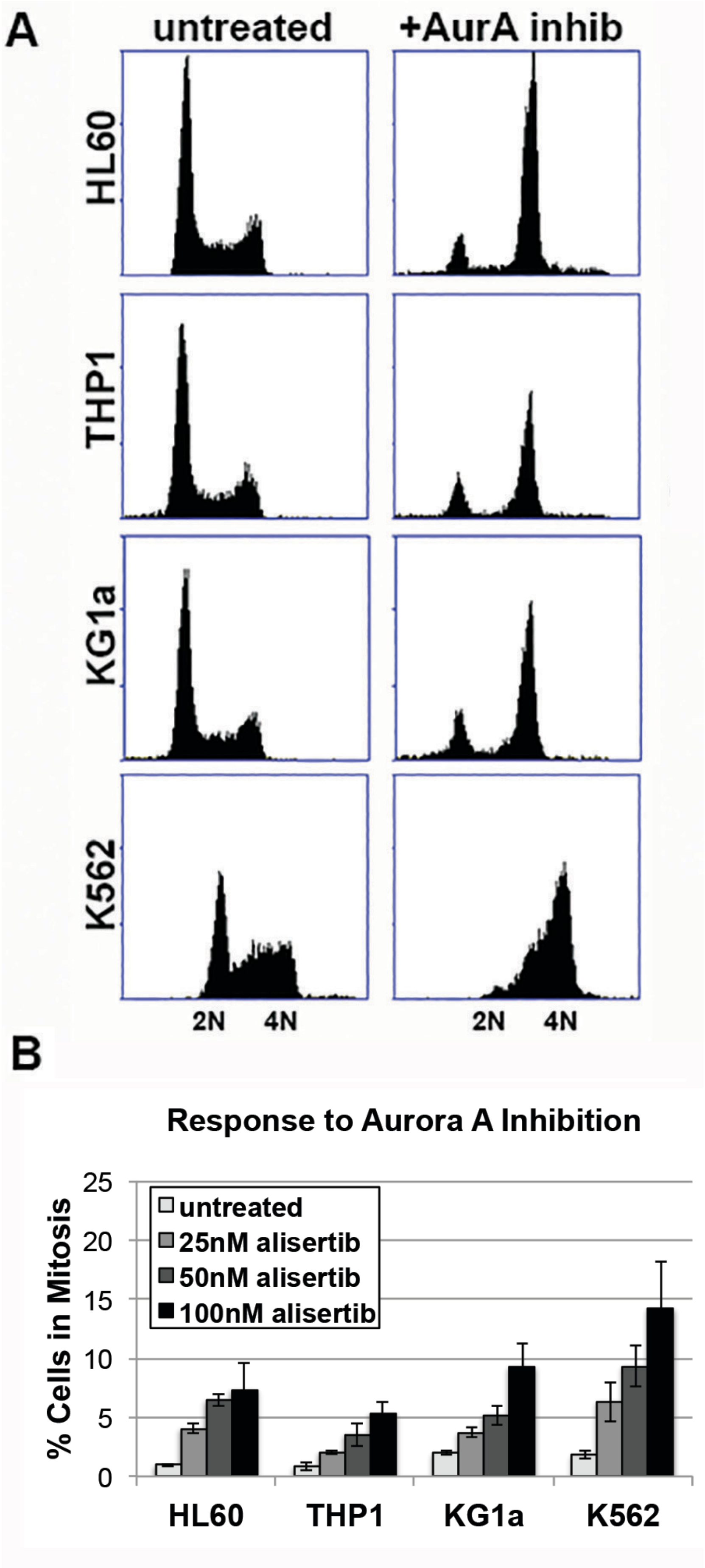
A
untreated
+AurA inhib
HL60
THP1
KG1a
K562
2N
4N
2N
4N
B
Response to Aurora A Inhibition
% Cells in Mitosis
25
20
15
10
5
0
untreated
25nM alisertib
50nM alisertib
100nM alisertib
HL60
THP1
KG1a
K562

Supplemental Figure 4

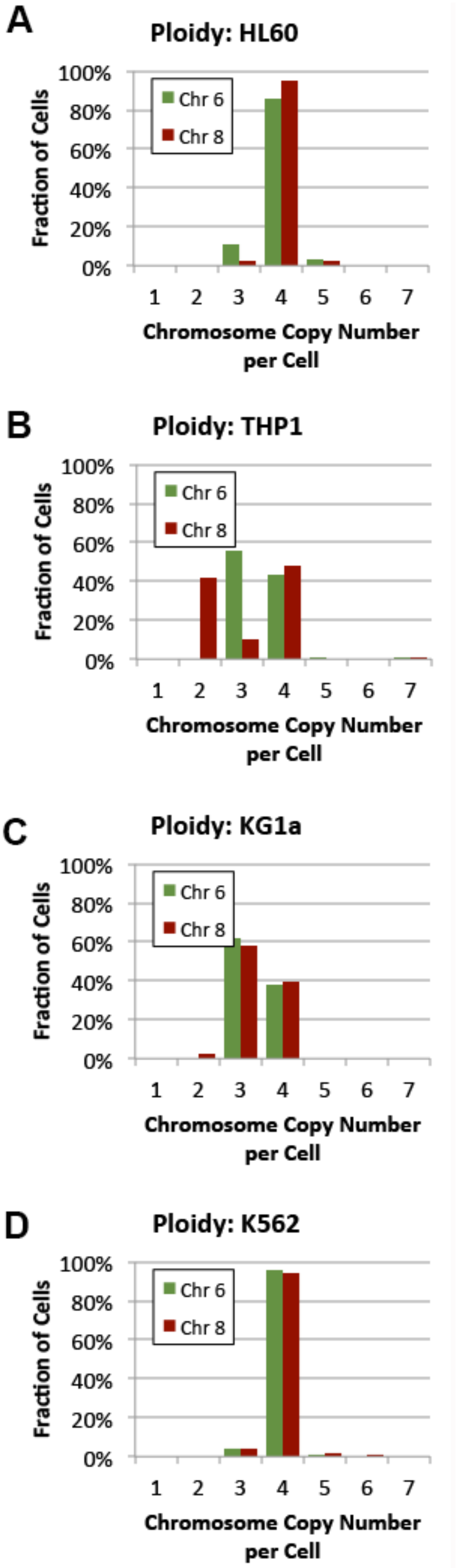
